# A structured reflective process supports student awareness of employability skills development in a science placement module

**DOI:** 10.1101/2021.02.07.430108

**Authors:** Luciane V. Mello, Tunde Varga-Atkins, Steve W. Edwards

## Abstract

Placements are often an extra-curricular activity of a science degree. This study reports on the outcomes of a final year credit-bearing 6-week placement module that was specifically designed to develop and enhance students’ employability skills. A key element of this module was that the student placements were not just evaluated from a science perspective, but with an emphasis on meaningful reflection and evaluation of employability skills development. Students recorded their levels of confidence in skills before, during and after the placement via an Online Reflective Log, as part of a module’s summative assessment. First, results showed that taking part in the placement and conducting their own independent research helped students to make connections between their scientific knowledge, otherwise constrained within the walls of the undergraduate science lab, and the wider impact of their research on society. Another theme that emerged concerned career choices and aspirations, and the placement experience either confirmed prior choices or opened new horizons. The Online Reflective Log helped students to feel supported by their university supervisor who were at a distance, while feedback on their tasks challenged them to reflect on the scientific and personal skills as they were engaged in scientific activities during placement. Students agreed that they had further developed their employability skills during the placement. Students acknowledged it was challenging to have to acquire evidence of skills development but appreciated the usefulness of this reflection in relation to their future careers.

## INTRODUCTION

In the 20 or so years since the Dearing Report [1], that identified the need to enhance a range of “key” competencies for university students to enhance their life- and employability-skills. Universities have responded accordingly, by either embedding these skills, including communication, numeracy, IT and learning skills, into core modules or by designing bespoke modules that develop such skills alongside the academic curriculum [2,3]. These changes have transformed skills development in students in spite of initial reservations by some staff that the University curriculum should only focus on academic subjects. The development of these skills had greatly aided student learning, but the focus has recently shifted to employability skills and preparedness of students for the workplace [4–6]. The inclusion of graduate employability statistics in the UK National Student Satisfaction (NSS) survey and the Teaching Excellence and Student Outcomes Framework (TEF) have highlighted the importance of this metric for Universities, and prompted further efforts to strategically embed employability skills more visibly in curricula [7]. The NSS is an annual survey for final year undergraduate students aimed to gather feedback on their experiences during their course; and the TEF assess excellence in learning and teaching and how Higher Education Institutions (HEI) ensures outcomes for their students in terms of graduate-level employment or further study.

Employers often report that they are generally satisfied with the academic abilities and standards of students and the value of their degree qualifications, but many still report that students are often not “workplace ready” and find it difficult to transition to a workplace environment from the academic environment [8–10]. This may be more marked when students seek employment beyond their academic discipline, and less apparent when students, for example, transition to study for a higher degree for which their previous academic experiences adequately prepare them. Work experience can help students prepare for the workplace, and both students and employers value the maturity, experience and understanding of the work environment that students gain during placements years [11–13]. However, the number of available year-long placements is declining and the national competition between students for such placements is fierce. Thus, not all students who seek a year-long placement and who would benefit from the experiences, are successful in obtaining these opportunities.

Many students do not fully recognise, reflect on, evidence and evaluate the full sets of skills that they have acquired during their University education and even if they do, they are often unable to articulate specific examples of how they can demonstrate competency in such skills at interview or in job applications [14]. Students are often assessment driven and unless employability skills are an integral part of the assessment process, they are often seen as optional, extra or co-curricular, activities. Students may not see the relevance of these skills, or more generally, how their University experiences prepare them for the workplace. However, skills development that is based on critical self-reflection [15,16] and experiential learning [17,18] can lead to greater student awareness and engagement, and so be more effective than the didactic teaching of such skills.

There are several definitions of employability skills, also known as transferable skills. The definition adopted in this study is that of Knight and Yorke [19] as ‘a set of achievements, understandings and personal-attributes that make individuals more likely to gain employment and be successful in their chosen occupations’. While *employment* should be regarded as an outcome, *employability* should be viewed as a lifelong attribute [20]. Lifelong learning involves understanding how to learn, reflect and progress [21]. It is founded on the premise that a greater awareness and understanding of ourselves can improve the way we work, contributing to our personal development, but also to society as a whole.

In this article we report on the outcomes of a 30-credit Master level (UK level 7) module that was specifically designed to develop and enhance employability skills of science students via a 6-week placement at a UK organisation or overseas-partner University. The purpose of this module is for the students to gain work-based experiences that cannot be obtained during conventional projects in our host laboratories. Key to the learning outcomes of this module are: (a) instructional guidance to the students on the expectations of the placement and their role in the experiential learning process and personal development; (b) self-evaluation by students of their perceived competencies in a range of employability skills before, during and after the placement, coupled with self-reflection of how these skills have been developed and can be demonstrated, and (c) awareness of the impact or beneficiaries of the work being undertaken and how this work can benefit society. External examiners and employers have commented very favourably on the outcomes of this module on students’ skills development and preparedness for the workplace. The aims of this study were (a) to determine the students’ perceptions of a proposed structured reflective process that supports the development of their transferable skills whilst on placement, (b) to explore students’ perceptions of the value of this module for their career development and employability prospects, and (c) to assess their awareness of the personal skills developed during a placement.

## BACKGROUND

### The module and placements

The placement module is for students on the Integrated Masters (MBiolSci) of the School of Life Sciences at the University of Liverpool, UK. An Integrated Masters in the UK is a four-year degree, where the first three years constitute the bachelor years, and the fourth year is the master year. The placement activity runs in the summer between years 3 and 4 of study and must be conducted at a location outside the School of Life Sciences.

There are many types of placement opportunities available to the students. They should select one that will give them the type of additional experience that will be important for their future career. Students have the opportunity to work abroad or in the UK. The idea is to experience a very different approach to research influenced by local factors (economic or social) that are pertinent to the region of the world in which they are working. For both types of placements, national and international, students should understand the economic or social factors influencing their work/research and should learn about *why* the particular work is being done and *who* is likely to benefit if the work is successful. During the placement, a staff member from the host institution supervises the student’s work. In addition, a University of Liverpool staff member provides the support needed for the student’s learning and personal development. This includes a combination of pastoral support and guidance during for students’ self-evaluation of the perceived competencies developed in a range of employability skills.

### Recording and evaluating skills

Based on the knowledge that bioscience students need more training and guidance in reflective practice [22] and students are assessment-driven [23], the Online Reflective Log (ORL) summative assessment was introduced to the module in the 2016-17 academic year and is completed before and during the 6-week placement. It is weighted as 25 % of the total module assessment, engaging and supporting students in reflecting on skill development [24]. The other two assessments, completed after students return to the university campus, are the placement report and an oral presentation, weighted as 65 % and 10 % respectively. The Online Reflective Log aims to foster students’ self-evaluation as students have to reflect on and record evidence of their acquired transferable skills. Students in the past tended to struggle when asked to reflect on their development and acquired skills. For this reason, their University supervisor offers feedback and guidance on the completion of the ORL to students during their placement. In addition, the Log helps the university supervisors to monitor and identify if students encounter any problems during their placement; they are then able to support students while off-campus. The Online Reflective Log has three main learning & teaching purposes: 1) formative feedback (guidance to the new placement experience encouraging students to evaluate their employability skills development); 2) summative assessment (provide a mark to the work as a way to boost engagement); and 3) diagnostic (a regular dialogue between university staff and students allow problems identification).

Students should engage with the Online Reflective Log prior to starting the placement, then every week during the placement. The Log is divided into three main parts as shown in Figure 1a and b. The first, the *Introduction & Goals* part, focuses on providing information on the module learning outcomes and assessments. In addition, it asks the students to provide information about the placement host institution, and the reasons for taking the placement – its goals and objectives. It is worth noting that these are not in relation to the placement research but rather what employability skills can be developed during this placement experience. The second part is the *Skills Audit* that consists of 13 skills and attributes. Students are asked to perform a self-assessment using a scale of 5 (1 poor – 5 outstanding) (Figure 1b). This exercise is done three times, pre-departure, and in weeks 3 and 6. In week 6, the last week of the placement, students are also asked to explain the scores given on the three occasions, providing evidence (250 words per skill attribute) for any change, or no change, in the score; reflecting on their personal skills and development. The third part is *the RPR (Recording, Planning and Reflection*). This consists of a brief description of the work done in the week, and two questions: (a) what the student learned/gained from the work done in the week (scientific and employability skills), and (b) the plans for the next week. Students complete this weekly activity in the Online Reflective Log by the end of every placement week.

**Figure 1.**
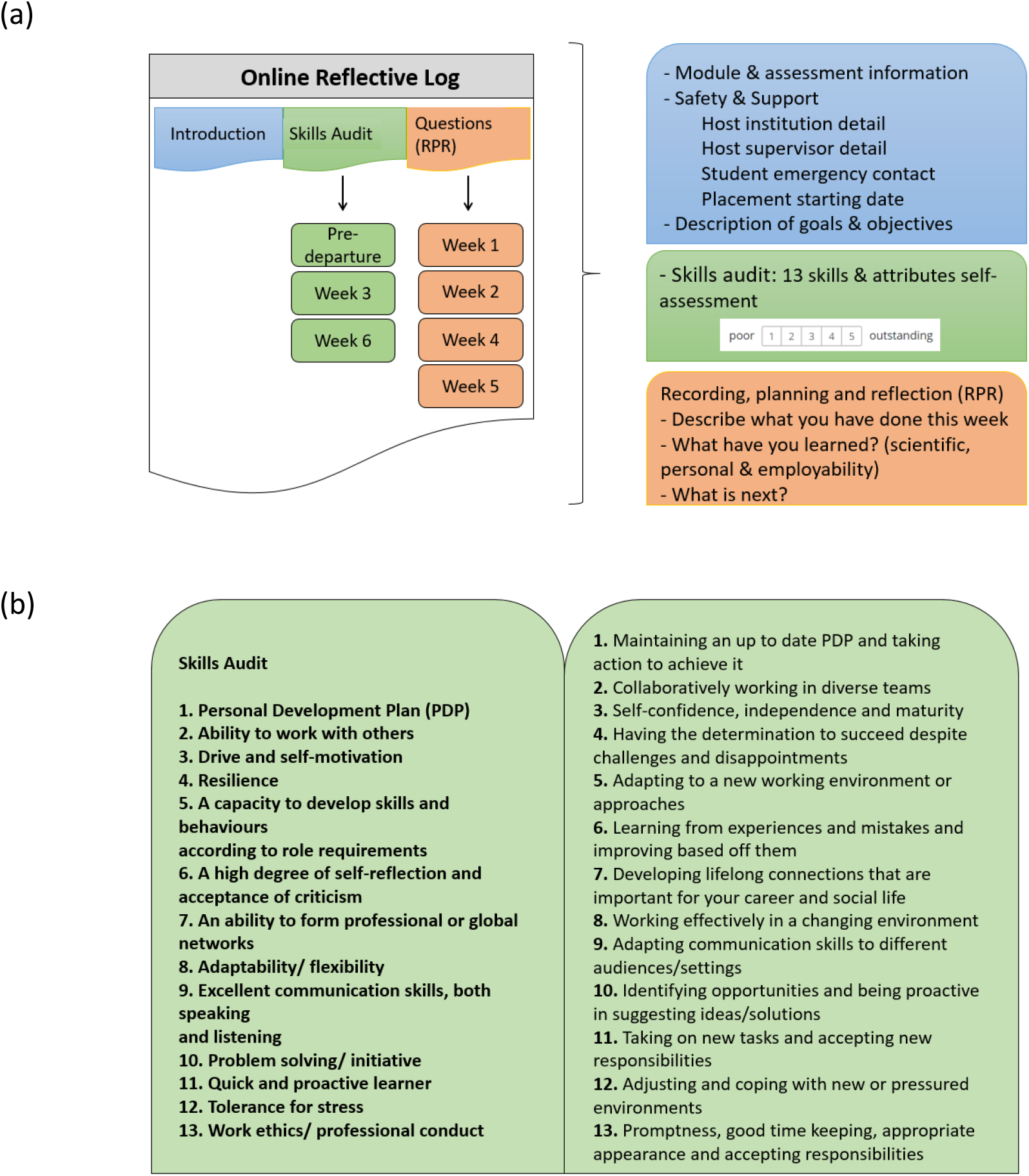
Online Reflective Log (ORL): (a) The three components of the ORL: introduction, skills audit and questions. The time of completion: pre-departure and last day of the indicated week in the boxes; (b) Skills Audit: students score themselves (pre-departure, week 3 and week 6) and give evidence on 13 main attributes and employability skills (week 6).

In summary, the six weekly online forms contain specific elements for students to complete, which are then read and evaluated online by University of Liverpool supervisors who provide timely feedback and suggestions for improvement. This log also includes a skills audit, highlighting both the scientific and personal skills that students have developed during their placement. Monitoring students’ ongoing performance is important in order to encourage formative feedback and dialogue between students and the University supervisors, as well as encouraging students to take responsibility for their learning.

### Research questions

This study’s research questions were: 1) What are students’ perceptions of the value of this placement experience for their career development and employability prospects? 2) How do the students recognise the skills developed as part of their placement? 3) What was the impact of the Online Reflective Log on students’ skills development?

## METHODS

Ethics approval was granted by the University of Liverpool’s Ethics Committee in March 2016. Informed consent was sought and granted by participants. Punch [25] describes empirical research as research based on direct experience or observation of the world involving two main kinds of data, quantitative or qualitative. A questionnaire and a focus group were used to collect student feedback after the introduction of the Online Reflective Log to the module. In addition, students were invited to comment on the module in a school-wide end-of-module evaluation. A structured focus group was facilitated to enable students to freely discuss the online resources, and encouraging them to think about their placement experience and how it was evaluated. Free-text comments from questionnaires as well as the focus group transcript were analysed using thematic analysis [26]. All participants were fourth year students in the School of Life Sciences. Students were provided with information sheets prior to their participation and participants signed consent forms for the focus group.

### Focus Group

A focus group is a way of conducting a collective interview, using an unstructured, semi-structured or structured approach [27]. The focus group took place with eight fourth-year students from the School of Life Sciences in 2016-17, facilitated by one of the authors (TVA). An email was sent out to all fourth-year students inviting them to participate in a focus group, which explored themes arising from a pilot survey that students had previously completed about their placement experience (not shown here). Participants were selected as they took part in an elective 6-week summer placement, abroad or in the UK. The host sites included research labs, research-support offices, clinical trial units and hospital settings. The 60-min session was recorded and transcribed. The session was structured exploring the following themes: placement experiences; impact of the placement; employability skills acquired during the placement; and skills development enhanced by the Online Reflective Log exercise/assessment. The focus group findings were fed into the design of a new questionnaire to evaluate this study.

### Questionnaires

Questionnaire-based research is a well-known and commonly used research approach in educational research [28]. This study used two different questionnaires to collect students’ views: the study questionnaire and the module evaluation survey used for evaluating modules in the department. Questionnaire data were collected for three consecutive academic years: 2017/18, 2018/19 and 2019/20. The number of students on the module for these academic years was N= 38, N= 23, N=31, N=30, respectively. Students completed both questionnaires voluntarily and anonymously after all the coursework and assessments.

The study questionnaire, designed based on the focus group analysis, contained ten questions; two consisted of Likert scale data exploring students’ perception of the impact of the placement on their skills development. Three were open questions asking the students to list the most and least beneficial aspects of taking a placement, and the impact of a placement on their future career plans. Two questions addressed the Online Reflective Log assessments as learning opportunities. The remaining three questions were: a demographic one, the students’ Master specialisation subject; and suggestions for improvement of the placement provision. In addition to this questionnaire, we analysed data collected via the module evaluation survey in all three years of study. This is a standard questionnaire for all modules, designed to gather students’ feedback on the module, including questions on content, teaching approach and assessments. For this study, we focus on the open questions of the evaluation questionnaire, where students are asked to list up to three particular strengths of the module.

## RESULTS

The quantitative and qualitative results were analysed in three main categories according to the aims of the study: students’ perception of (a) the impact of the Online Reflective Log on their skills development; (b) the value students placed on their placement experience for their career development and employability prospects; and (c) students’ perception on the skills developed as part of their placement. Quantitative questionnaire data are mentioned as three values, corresponding to the three academic years of data collection. The response rates of the study questionnaire were 82 %, 78 % and 74 % in the academic years of 2017/18, 2018/19, and 2019/20, respectively. Figure 2 shows that students almost always strongly agree or agree that the placement experience benefited their skills development, and that the introduction of the Online Reflective Log assessment to the module was essential to support the reflective exercise requested from the students in this module.

**Figure 2.**
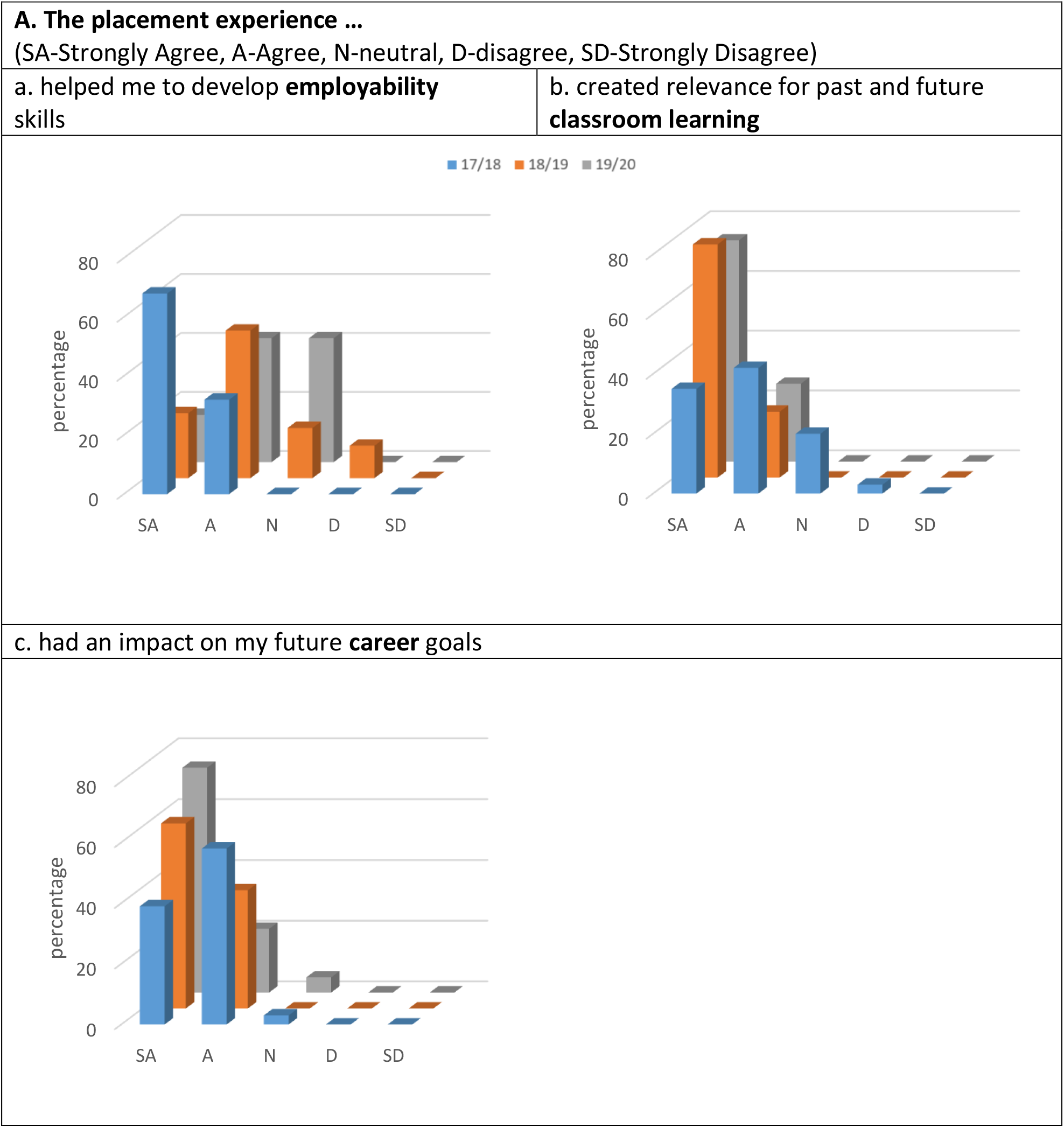
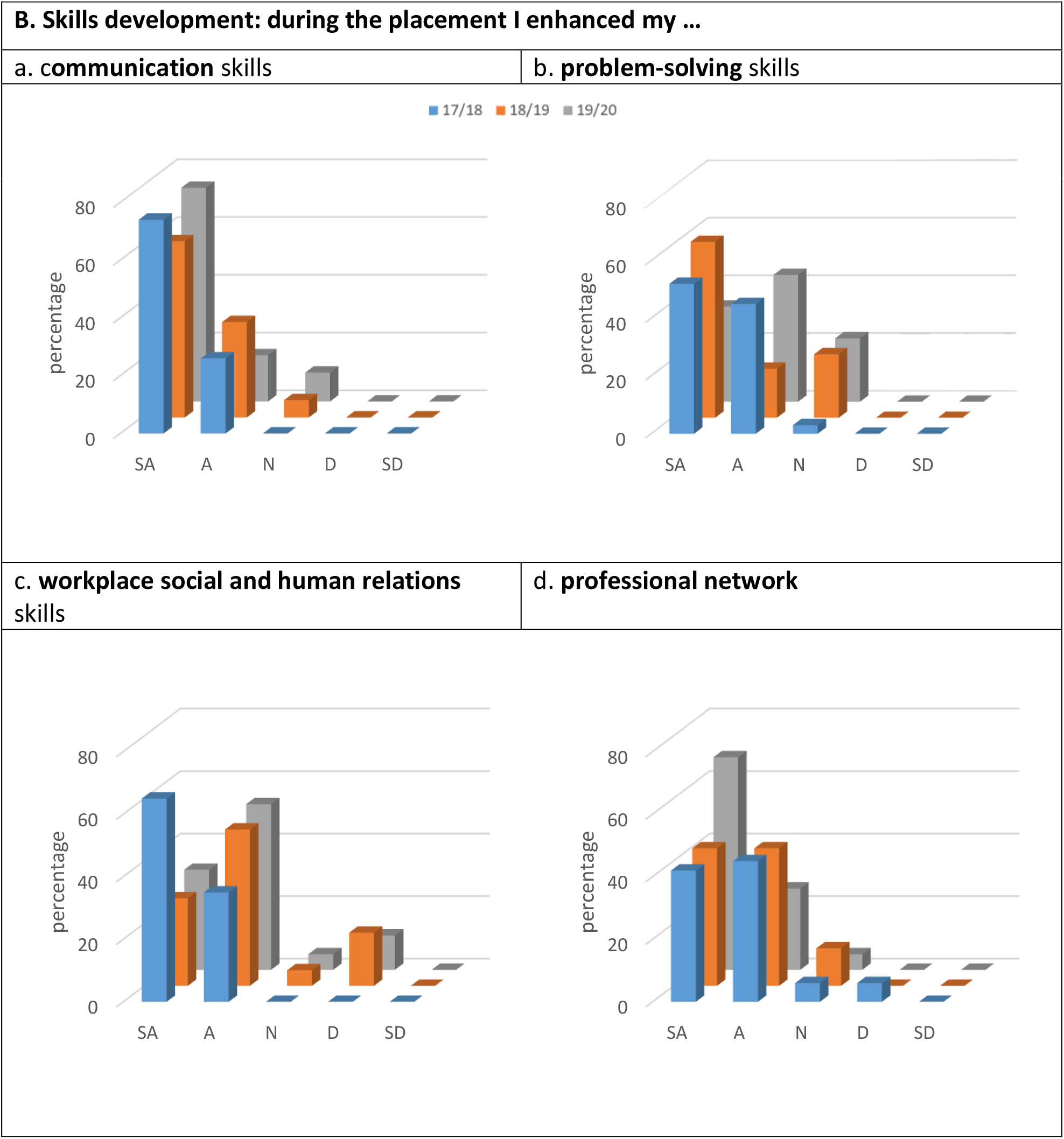
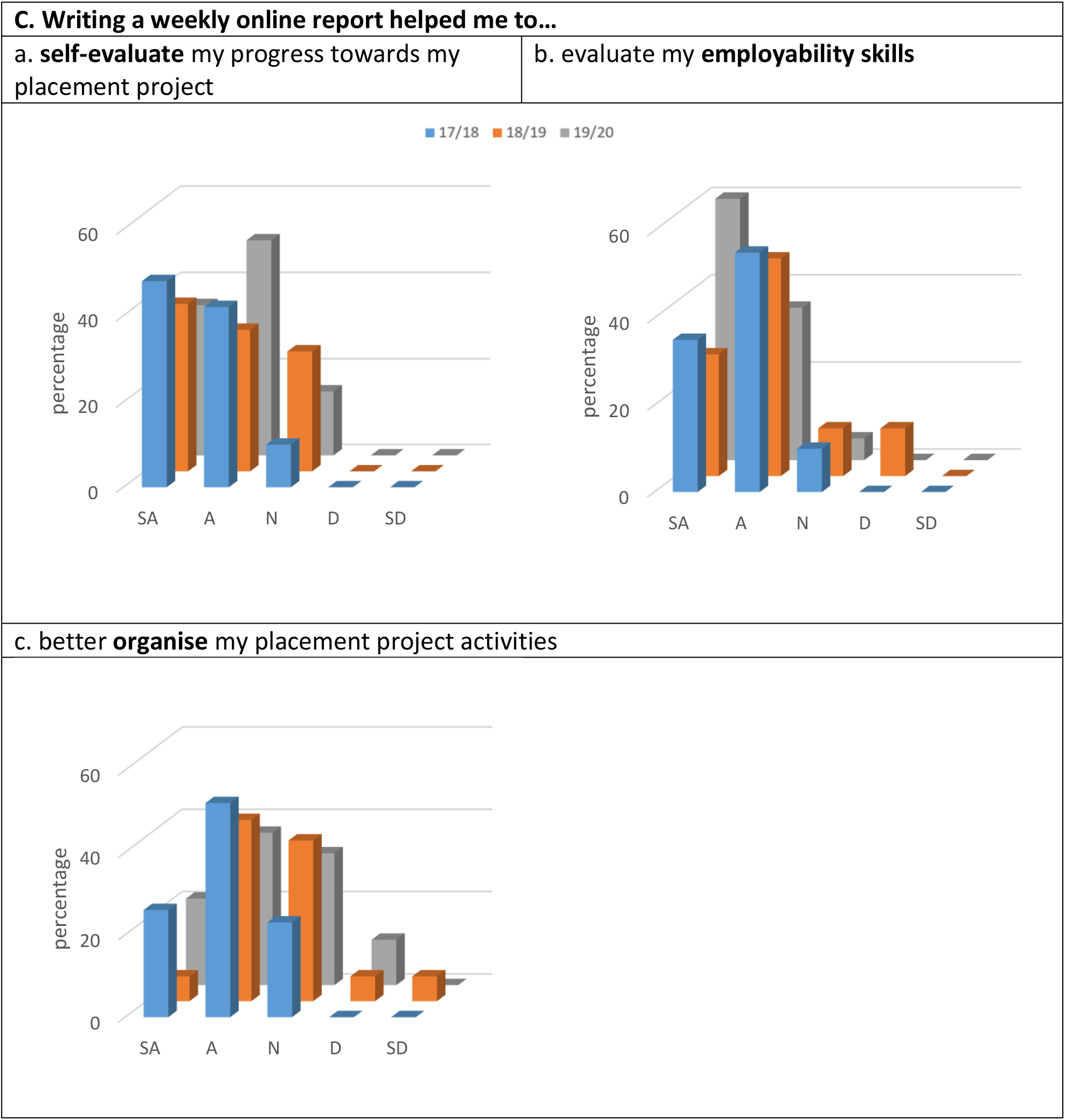
Percentage of respondents agreeing/disagreeing with the several statements on skills development during the placement. Cohort size and % of respondent for each year of analysis: 2017/18, N=38, respondents=82 %; 2018/19, N=23, respondents=78 %; and 2019/20, N=30, respondents=74 %.

**Figure 3.**
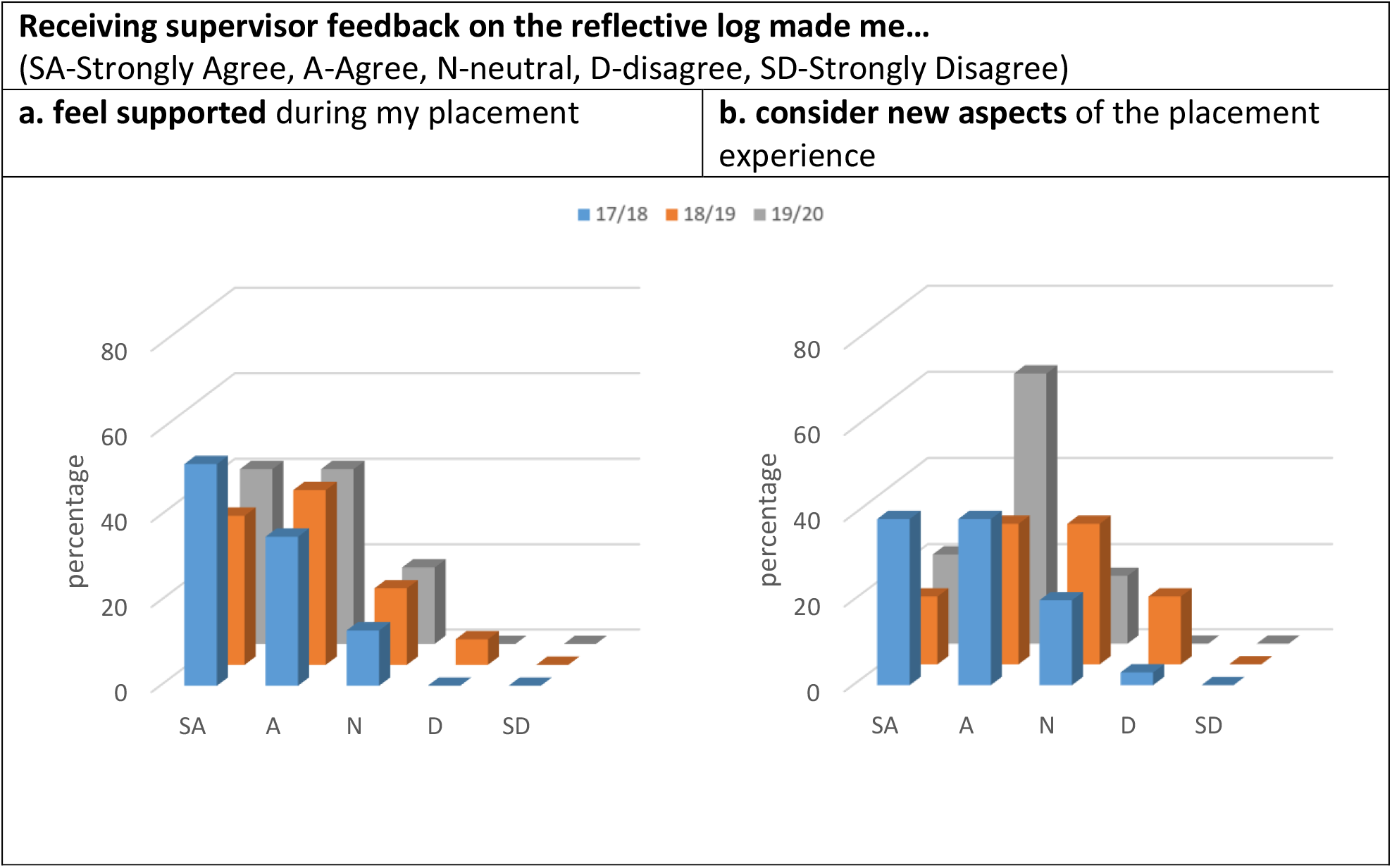
Percentage of respondents agreeing/disagreeing with the statements on receiving feedback on the online reflective log during the placement. Cohort size and % of respondent for each year of analysis: 2017/18, N=38, respondents=82 %; 2018/19, N=23, respondents=78 %; and 2019/20, N=30, respondents=74 %.

### The placement experience

Students were positive about their placement experience. Figure 2(A) shows that 84%, 72 % and 58 % of the students either strongly agree or agree that the placement experience helped them to develop employability skills; 77%, 100% and 100% either strongly agree or agree that the placement experience created relevance for past and future classroom learning; and 97%, 100% and 95% either strongly agree or agree that the placement experience had an impact on their future career goals. These results were supported by the qualitative data of the module evaluation survey that illustrates the most common benefits of the module mentioned by the students (Table 1). When compared to the focus group results, the findings reinforce the two themes that emerged with respect to the overall impact of the placement: linking the lab/work day-by-day with personal experiences, and expanding or confirming career aspirations. A number of students commented that, as a result of taking part in the placement and conducting their own independent research at the host site, they were better able to make connections between their scientific knowledge, usually constrained within the walls of the undergraduate science lab, and the wider impact of their research on society and people. One student commented that in the lab as a student they felt “*detached from any sort of benefit to people!*”, whilst during the placement they saw professionals in action, helping them to draw the links between science-in-the-lab and science as it impacts on people and society. Students related this realisation with the online reflective log (ORL) exercise, where they were constantly asked to reflect and discuss the purpose of their research project in line with the module aims (see results and analysis below).

**Table 1:**
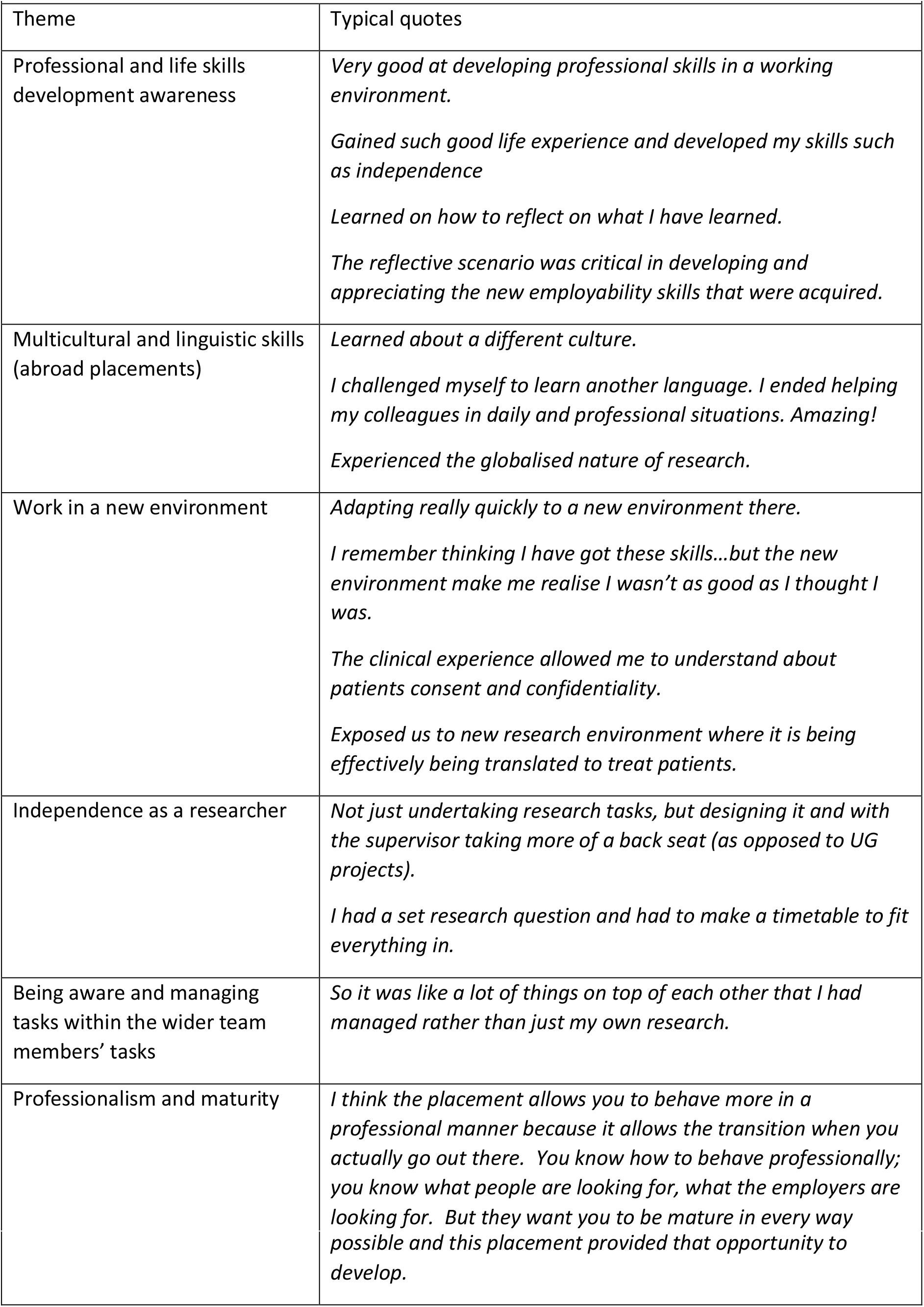
Themes and typical students’ comments on employability skills development during the placement.

Expanding on career choices and aspirations, the focus group showed that some students already had an idea of what setting they wanted to work in (e.g. pure research or clinical). However, they claimed that the placement experience had either confirmed this prior choice or opened new horizons. They were also able to appreciate a wider range of research settings (e.g. research-support or mix of clinical and laboratory research), of which they were previously not aware. In one case, the placement helped student re-assess their previous perceptions of certain research-related jobs, having earlier dismissed them as ‘office jobs’, but now seeing them as something more exciting and desirable. Others realised that they would like to have more links with the human/patient side of the research and started considering a clinical setting for their future career. This diverse response reveals the benefit of offering different types of placement, not only the most common industrial placements.

### The skills developed during the placement

Students were extremely positive about their development of the four skills described in Figure 2B with responses of either ‘strongly agree’ or ‘agree’ of: 100 %, 94 % and 90 % for communication skills; 97 %, 78 % and 77 % for problem-solving; 100 %, 78 % and 85 % for social and human relations; and 88 %, 88 % and 95% for professional networking development. It is worth noting that for the academic year 2019/20 this result contradicts those from Figure 2A(a), where a high number of students said the placement experience was neutral with respect to the development of their employability skills. A similar positive response was seen in the quantitative analysis of the final year module evaluation, where students saw the link between communication skills, human relations and networking (Table 1). In the focus group, students expressed their appreciation of being in a new setting for their research experience, as they appreciated the learning process in relation to networking with colleagues: “*I would say I learnt the most from actually talking to the researchers and sort of just working within the team*”. Table 1 shows further examples from the focus group discussion of students developing employability skills as a result of the placement. One of the students commented that although they were actively looking through job applications during their degree and learning what employers are looking for via the job specification, it was only through the placement that he was able to understand what these skills meant. Many students also mentioned this realisation in the final year module evaluation, linked to the importance of the Online Reflective Log: “*the placement taught new skills and the reflective scenario imposed by the reflective log was critical in developing and appreciating the new employability skills that were acquired*”.

### The impact of the Online Reflective Log on students’ skills development

The Online Reflective Log consisted of three main parts: introduction, skills audit and questions. The introduction part was used for information about the student and the placement. This study analyses students’ responses to the last two parts, the skills audit and questions (Figure 1).

Figure 2(C) shows the student’s evaluation of the benefits of the Online Reflective Log prior to and during their placements. Once again, students almost always ‘strongly agreed’ or ‘agreed’ with the statements: 90 %, 72 % and 85 % for self-evaluation of students’ progress towards their placement project; 90 %, 78 % and 95 % for evaluation of their employability skills; and 81%, 83% and 84% for organisation of their placement project activities. However, for this last category, a higher proportion of students chose ‘neither’, indicating a less beneficial impact of the Online Reflective Log for this particular activity. As previously mentioned, one of the aims of the Online Reflective Log was to improve support to the students while off-campus by identifying their development needs. When asked about the benefits of receiving regular feedback from their academic supervisor via the Online Reflective Log, 87 %, 76 %, and 82 % of the respondents either ‘strongly agreed’ or ‘agreed’ that they felt supported during their placements. In addition, 78 %, 49 % and 84 % revealed that the feedback made them consider aspects of the placement experience that they had not thought about before their interaction with the University of Liverpool supervisor (data not shown). The Online Reflective Log was praised as an excellent learning tool, regularly mentioned in the final module evaluation survey and focus group. A student commented: “*The weekly Online Reflective Log was great for keeping students motivated and forced students to think about the cultural and employability aspects*”. While another student said that the Online Reflective Log “*challenges you to reflect on your personal development*”. The comments from the focus group were grouped by themes and are presented in Table 2.

**Table 2.** Students’ comments on the use of the Online Reflective Log (ORL) during the focus group.

| **Table 2. Students' comments on the use of the Online Reflective Log (ORL) during the focus group.** |
| --- |
| **(a) The weekly ORL helped you remember:** |
| *It was really useful to have the reflective work that we'd done every week because then you could just look and see what you've been doing and just add it all together and decide. So it was all there so you haven't forgotten anything like you might have if you weren't logging it down every week.* |
| *It is only for 6 weeks: timeframe awareness.* |
| **(b) The impact of taking notes:** |
| *It is the writing down process that makes you realise what you have developed, or picked something up.* |
| *When you write it down you realise you've actually picked up something and that's one of the skills the employability skills that the employers look for.* |
| **(c) Reflection:** |
| *I am not a reflective person – the skill to reflect didn't come naturally to us, it was something that we had to learn, which in turn helped us to be able to self-evaluate our actions.* |
| *Reflection does not come naturally and I have never asked to reflect on my development, at this extent, before.* |
| **(d) Progression and development – whether as scientists, researchers or professionals:** |
| *As the 6-weeks passed, looking back through the logs, we could see that we could do more work in the time allotted.* |
| *The frequency of this task made us more aware of their timeframe. It encouraged us to do more with your time....critique yourself as well.* |
| *The audit skills is a good opportunity to evaluate yourself.* |
| **(e) Use of literature in employability:** A particularly useful aspect of the ORL was that students were encouraged to consult the literature on what employers were asking; this helped them be more aware of what skills they were developing: |
| *It's not necessarily picking up the skills but looking through the literature which they kept encouraging us to do. It did actually make you realise some things that employers might be looking for that you weren't aware of. Like one of the things that I found was quite a large percentage wanted employees to be open to working in different environments and maybe across the globe and that experience of working abroad like made me want to work in different countries throughout my career.* |

A particular area explored during the focus group was the reasons for fluctuation of self-audit scores - students’ reflection on the skills audit scores (Figure 1). Some students reported a fluctuating trend (higher-lower-higher scores) for their self-evaluation of skills pre-during-post placement, for instance: “*This sounds really arrogant but I remember thinking I’ve got some of these skills and I think I’m going to be quite good at this and then by week three, because you’ve been up half the time and you’ve got used to it, you realise that maybe in that environment because it was new you weren’t as good as you thought you were… so that brought my scores back down and towards the end the other thing was coming to a close and starting to get all the results, it brought my scores back up because I was like I’ve managed to achieve all of this and I’ve done all of these things that maybe week three me would’ve struggled with. So I think that’s why mine [came] back up in the end*”. Another student highlighted the helpfulness of being supported to audit and reflect on their skills via the weekly logs: “*I actually thought it was really helpful doing the audit because it did let me put into context what I’d been doing and the fact that my academic supervisor asked us to write reference papers and that meant I had to go out and read things. So I did find it quite helpful but I started out thinking my professionalism, my time management is great and then midway through you’re okay it could be better like when you compare it to people in a professional setting rather than in my honours project setting*”.

As the intended outcome of the reflective process was for students to recognise and evidence their skills, such fluctuation in reported skill levels was expected by university supervisors. However, as the placement progressed, decreasing scores were perceived as indicative of failure by some students. This showed that a clearer explanation of the aims of this exercise is needed, namely to: increase awareness of personal strengths and weaknesses, followed by a plan for further development of each skill. This observation led to changes in the module, and from the academic year 2018-19 more support for skills audit completion was offered to the students, prior departing to their placements.

## DISCUSSION AND CONCLUSION

The aims of this study were to determine: (a) the impact of undertaking a reflective process on students’ perceptions of their skills development through the Online Reflective Log summative assessment; and (b) the value students placed on their placement experience/module for their career development and employability prospects; and (c) students’ perception of the skills developed as part of their placement. To meet the study aims, we gathered students’ views using a focus group and surveyed the students for three consecutive years, minimising any cohort bias.

Our results show that the placements helped students to develop both scientific and transferable skills, and one may argue this is a well-known outcome. However, this study shows skills development during a six-week placement, instead of the more conventional year in industry placements. Furthermore, the reflective process structured by the Online Reflective Log assessment was key to raising students’ awareness of their strengths and weaknesses, supporting life-long learning. The benefits of using reflective logs to support students’ development of employability skills have already been established [29]. More recently, Hill and collaborators [30] showed that science students reported ‘unfamiliarity of thinking beyond knowledge attainment in order to identify and reflect on skill-related experiences’. Their study also showed that students recognised that by improving their ability to reflect, they would be better prepared to articulate their skills when seeking employment. The need to improve students’ reflective ability was also illustrated by Voelkel and collaborators [31], where it was shown that students’ willingness and ability to reflect on their learning process during their final year dissertation was patchy, indicating that more training in the skills of reflection earlier on in their course was needed.

Building on these findings, we can say that one of the main contributions of this paper to the literature on placements and skills development is the focus on the ongoing formative feedback and summative assessment of skills development, including students’ self-reflection and recording of their development, as part of a credit bearing module. Placement students tend only to be assessed through a report on their placement work and experiments i.e. on their scientific results. In this study, however, we encouraged the students to reflect on the process by which they gained and developed employability skills, and to use this reflective process to design their own personal development plans. While other studies have reported on the use of skills audits to help students’ self-evaluation of acquired skills in response to a particular activity, the reflective component has not been explored [2,32]. Kensington-Miller and collaborators [33] presented a framework to help undergraduates recognise, articulate and evidence attributes or skills that are often developed or required within a university degree and profession, but are not explicitly discussed or assessed. In this module, students have to explain the changes in their skills audit scores, e.g. they need to provide evidence on how skills were developed, or seemed to be lacking*: “ it is the writing down process that makes you realise what you have developed, or picked something up*” (student).

Beard [32] claims that “*the value of a placement experience is enhanced when the experience is systematically assessed*”. The results of our study showed the benefit of reflective activity embedded in a credit-bearing module to foster skills development. The Online Reflective Log helped students to critically assess their skills and keep a permanent record of performance and development. We argue that recording skills may serve as a launchpad for continuing professional development. Studies show that employers often look for ‘soft’ skills as critical for professional success alongside traditional ‘hard’ skills [11,34]. Thus, it could be argued that the structured reflective process as part of the Online Reflective Log should better prepare the students for future job applications and interviews, increasing their employment chances, whether in industry, in clinical settings or in academia.

The science curriculum should allow for more reflection and employability skills development. A common trend observed by supervisors when providing feedback on students’ Online Reflective Log tasks during their placement (Recording, planning and reflection section) was that in the first two weeks, students mainly focused on reporting their weekly scientific lab experiments: what was done and plans for the following week. The feedback from the university supervisors was essential to bring additional transferable skills development experiences and planning into students’ discussions. This demonstrates that skills development reflection does not come naturally to students studying science. The reflective process should go beyond an organised and detailed lab book: it should go beyond scientific skills. Therefore, we emphasise the importance of the reflection process being well structured and supported by university supervisors. This means paying attention to: (a) How often do students evaluate their skills development; and (b) How do we help students with skills recording and reflection.

Another aspect to consider is that in general, placements either are offered in Higher Education Institutions as extra-curricular activities, or have a low weight towards a student degree; e.g. a year in industry may count for only 10 % of the overall degree mark. Other studies present ideas of how to develop extra-curricular activities to develop students’ employability skills (e.g. [2]. The module described here is worth a total 25 % of the students’ Master year total credits (30 out of 120 credits), demonstrating our emphasis on skills development in the curriculum. In addition, the Online Reflective Log here proposed was used as a summative assessment weighted 25 % of the module. The summative aspect was used to ensure students’ engagement with the reflective activity. This module has developed further in the current academic year, 2020-21. We are now working in collaboration with the university’s Careers and Employability Centre who supplement the placement pre-departure activities by delivering three talks: what employers look for in graduates; what you can gain from placements; and how to complete a reflective log. The benefit of the Online Reflective Log on students’ learning can be demonstrated by its incorporation into three other modules in the School of Life Sciences. Students are now asked to complete a skills audit in years 1, 2 and 3, and reflect on their development during their years of study. The Online Reflective Log has been adopted by other Schools/Departments at our university, and its benefits for supporting students off-campus can be illustrated by a quote from a colleague: *The strategic Online Reflective Log monitoring system is a powerful tool to support students on placement. It allows us to check on the students’ wellbeing while off-campus, and guide employability skills development*. Outside our institution, the Online Reflective Log with the structured support by academic supervisors has been implemented as part of other placement activities/modules: a colleague at a different university reported that they adopted the Online Reflective Log in the 2019-20 academic year. Furthermore, they noted that students were writing reflective diary style notes in the margin of the lab books as their project progressed. This demonstrates the importance of establishing a structured process of supporting students towards being able to recognise the transferable skills they are developing as part of their degree course bridging the gap between the classroom and the workplace.

Another unique aspect of this study was to focus on the outcomes of a six-week science placement instead of a year-long one. Students’ responses demonstrated evidence that skills development can be achieved during a shorter placement. In addition, the placements were offered to students in several settings, not just in industry, as students’ careers aspirations vary. The results illustrate that different working environments provide the students with common skills development, while they can also develop students’ appreciation of different challenges; e.g. cultural and language barriers, and different priorities in industrial, clinical and academic settings [35].

In summary, the module was well received by the students and has allowed them to develop key employability skills within a shorter, six-week period. The Online Reflective Log has emphasised students’ initial difficulties with the reflection process, but has also demonstrated that if properly supported and supervised, students will engage with the reflection. There is no doubt that a placement experience fosters skills development, but a well organised and carefully structured, supervised and assessed experience can help students to achieve a better understanding and awareness of their skills development. Placement assessments should not only focus on the final scientific outcome but on the whole process of personal and scientific development.

## Acknowledgements

This work was funded by the School of Life Sciences, University of Liverpool, Liverpool, L69 7ZB.

## Author contributions

LVM conceived, and LVM and SWE designed the project. LVM and TVA acquired, analysed and interpreted the data. LVM and SWE wrote the manuscript. TVA commented on multiple versions of the manuscript.

